# Testing Rare-Variant Association without Calling Genotypes Allows for Systematic Differences in Sequencing between Cases and Controls

**DOI:** 10.1101/032037

**Authors:** Yi-Juan Hu, Peizhou Liao, H. Richard Johnston, Andrew S. Allen, Glen A. Satten

## Abstract

Next-generation sequencing of DNA provides an unprecedented opportunity to discover rare genetic variants associated with complex diseases and traits. However, when testing the association between rare variants and traits of interest, the current practice of first calling underlying genotypes and then treating the called values as known is prone to false positive findings, especially when genotyping errors are systematically different between cases and controls. This happens whenever cases and controls are sequenced at different depths or on different platforms. In this article, we provide a likelihood-based approach to testing rare variant associations that directly models sequencing reads without calling genotypes. We consider the (weighted) burden test statistic, which is the (weighted) sum of the score statistic for assessing effects of individual variants on the trait of interest. Because variant locations are unknown, we develop a simple, computationally efficient screening algorithm to estimate the loci that are variants. Because our burden statistic may not have mean zero after screening, we develop a novel bootstrap procedure for assessing the significance of the burden statistic. We demonstrate through extensive simulation studies that the proposed tests are robust to a wide range of differential sequencing qualities between cases and controls, and are at least as powerful as the standard genotype calling approach when the latter controls type I error. An application to the UK10K data reveals novel rare variants in gene *BTBD18* associated with childhood onset obesity. The relevant software is freely available.

## Introduction

Recent technological advances in next-generation sequencing (NGS) have made it possible to conduct association studies on rare variants, which hold great potential to explain the missing heritability of complex traits and diseases (Manolio et al. 2009). However, it is prohibitively expensive to conduct high-depth, whole-genome sequencing (WGS) for large-scale association studies (Sims et al. 2014). Therefore, many WGS studies have reduced the overall average depth to as low as 4–10× (The 1000 Genomes Project Consortium 2012; The UK10K Consortium 2015; Morrison et al. 2013; Bizon et al. 2014). Other studies have adopted whole-exome sequencing (WES), in which only the protein coding regions were sequenced but at high depth (e.g., ≥30×) (The 1000 Genomes Project Consortium 2012; Tennessen et al. 2012; Epi4K Consortium, Epilepsy Phenome/Genome Project et al. 2013; The UK10K Consortium 2015); nevertheless, even though the *average* depth may be high, the large variability in capture efficiency may cause some genes or some regions within a gene to have much lower depth than the average (Do et al. 2012).

The case-control design remains the most commonly used approach to studying rare variant associations. Due to the high cost of sequencing, many studies have focused sequencing effort on cases. Some studies sequenced cases at higher depth than controls by design, when the cases are unique and there is interest in identifying novel mutations. An example is the UK10K Project (The UK10K Consortium 2015), which sequenced cases at ~60× and controls at ~6 ×. Some studies even sampled only cases for sequencing and intended to compare them with publicly available NGS data on general populations such as the 1000 Genomes (The 1000 Genomes Project Consortium 2012). In this case, the controls typically have systematically different sequencing qualities (e.g., depth and base-calling error rate) from the cases. Even when their *average* depths are similar, the *actual* depth could vary in individual regions across platforms, resulting in regions with differential depths in cases and controls by chance. This can easily occur when using different exome capture kits for cases and controls; if one kit can capture a certain exonic region better than the other, then there will be a systematic difference in read depth between cases and controls in this region.

The prevailing practice of analyzing NGS data for association with rare single-nucleotide variants (SNVs) is to first call underlying genotypes (e.g., using SAMtools (Li et al. 2009) or GATK (DePristo et al. 2011)), and then treat the called values as known in gene- or region-based tests such as the burden test (Morgenthaler and Thilly 2007; Li and Leal 2008). Genotype calling is difficult when read depth is low because minor allele reads are indistinguishable from sequencing errors. Genotype calling is especially challenging for rare SNVs, first because their locations cannot be easily inferred (Johnston et al. 2015), and second because little information can be borrowed from other variants through linkage disequilibrium (LD) (The 1000 Genomes Project Consortium 2012). In case-control studies with differential sequencing qualities, the genotype calling process can introduce confounding that causes inflated type I error in downstream association tests (Mayer-Jochimsen et al. 2013). Recall that confounding occurs when a variable is correlated with both the case-control status and the genotype. When read depths are different in cases and controls, the dependence of genotyping quality on the depth establishes the depth as a confounder. Likewise, the base-calling error rate has the same confounding effect as the depth. Even when read depths and error rates are comparable between cases and controls, differences in genotype calling algorithms or quality control (QC) filters (e.g., *phred* score cutoffs) can lead to differential genotyping errors that could also act as a confounder. For these reasons, publicly available NGS data have generally been under-utilized as controls for association studies. To reduce genotyping errors, one typically applies QC procedures to filter out SNVs at which many samples are covered by low depth of reads or called with low quality scores (Tennessen et al. 2012). The use of any reasonable QC procedure will remove a large number of variants, especially rare ones, and results in loss of important information.

To avoid the confounding effect induced by calling genotypes, Derkach et al. (2014) proposed to replace the genotypes in the standard score statistic by their expected values given observed read data, and developed a robust variance for the score statistic to account for differential variances of the expected values in high-and low-depth samples. However, they still used called genotypes to determine SNV locations, which tends to yield more false positive SNVs among the low-depth group (e.g., controls) than the high-depth group and again cause confounding. To ensure accuracy of the called SNV locations, they resorted to stringent QC procedures, which would result in substantial information loss.

In this article, we provide a likelihood-based approach to testing rare variant associations that directly models sequencing reads without calling genotypes. We consider the (weighted) burden test statistic, which is the (weighted) sum of the score statistic for assessing effects of individual variants on the trait of interest. Our read-centric approach enables us to explicitly account for sequencing differences (i.e., read depth and error rate) between cases and controls.

Full implementation of a read-centric approach requires solutions to a number of problems. Because SNV locations are unknown, we first develop a simple, computationally efficient screening algorithm to estimate their locations using read data alone. Because an imbalance in putative SNVs can arise due to differences in read depths and error rates between cases and controls, the burden statistic may not have mean zero even in the absence of association. Thus, we develop a novel bootstrap procedure for assessing the significance of the burden statistic. Specifically, we propose to generate a dataset with the same coverage patterns as the original data, but where the loci are all monomorphic. By comparing the false-positive SNVs found in the monomorphic dataset to the SNVs detected in the original data, we show how to estimate the number of true SNVs and the allele frequencies of the true SNVs in the original data. With this information, we can generate a bootstrap replicate dataset in which the allele frequencies at true SNVs match those in the original data, but are identical in cases and controls. We then compare the burden statistic from the original data to those from the bootstrap replicates to assess significance. The complete flowchart is depicted in Figure 1. Our procedure can encompass all informative loci including singletons and doubletons if desired; additionally, we can down-weight or mask loci that are unlikely to be deleterious.

**Figure 1.**
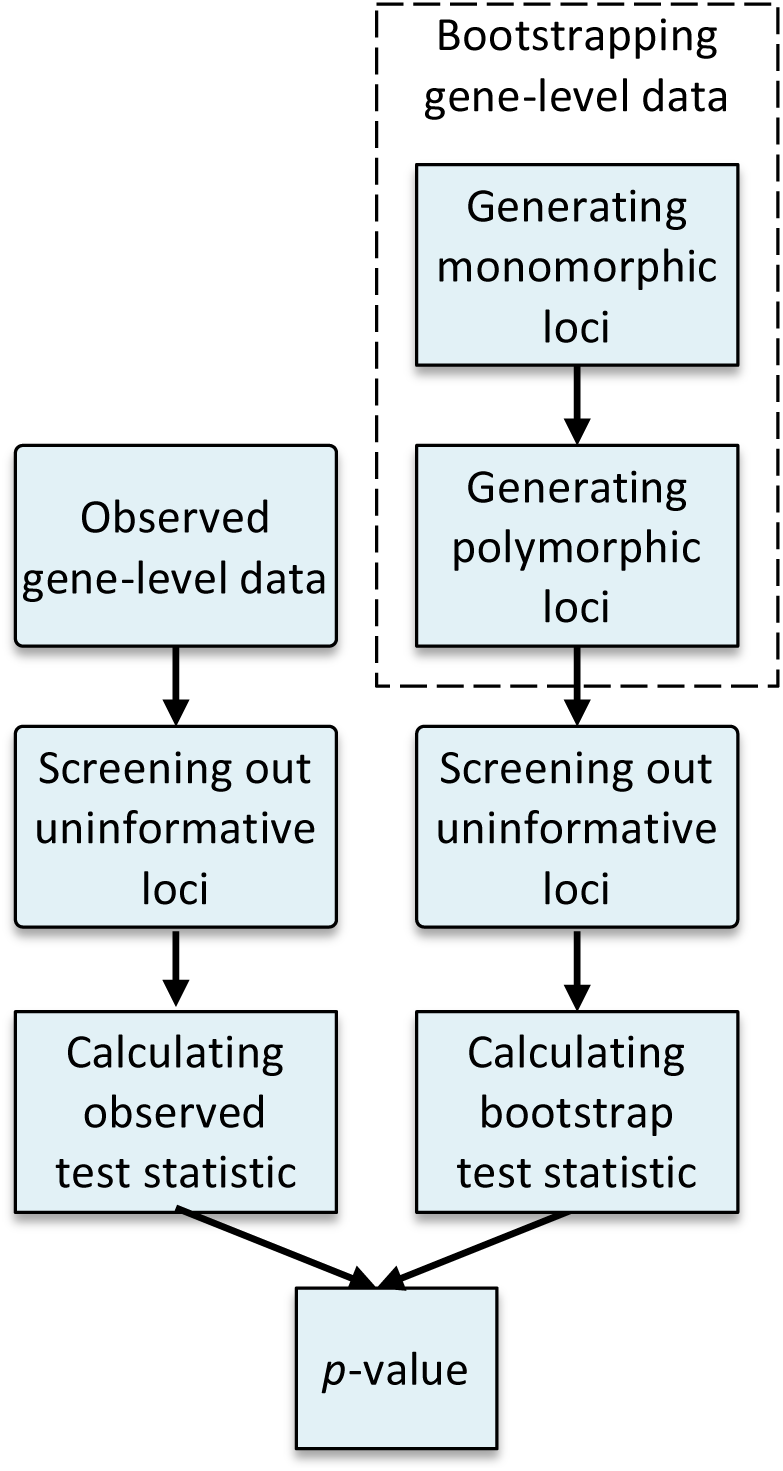
Flowchart of the proposed approach.

We showed through extensive simulation studies that our bootstrap tests are robust to a wide range of differential sequencing qualities between cases and controls, and are at least as powerful as the standard genotype calling approach when the latter controls type I error. We further applied the new methodology to a case-control data from the UK10K Project comparing children with severe early onset obesity to population-based controls. We identified a gene, *BTBD18*, that passes the exome-wide significance threshold and that is also a plausible candidate for childhood onset obesity.

## Methods

We first consider a single (bi-allelic) SNV. Let *G* be the genotype (coded as the number of minor alleles) at the variant site and let *D* be the disease status. We denote the genotype distribution under Hardy-Weinberg equilibrium (HWE) by *P_π_* (*G*), where *π* is the minor allele frequency (MAF). Note that the HWE assumption has a minimal effect for rare variants, as homozygotes of minor alleles are not expected. Instead of observing *G*, we observe the total number of reads mapped to the SNV and the number of reads carrying the minor allele, denoted by *T* and *R*, respectively. Similar to SAMtools, GATK, and seqEM (Martin et al. 2010), we assume that *R* given *T* and *G* follows a binomial distribution

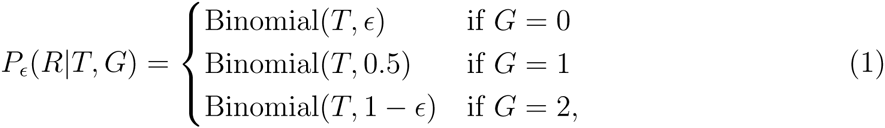

where *ϵ* is the probability that a read allele is different from the true allele and is referred to as the error rate. The “errors” here comprise both base-calling and alignment errors. We treat *ϵ* as a free parameter that is locus-specific and will be estimated from the read data (Martin et al. 2010).

### Test statistic

To account for case-control sampling, we use the retrospective likelihood 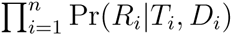 based on *n* subjects, which takes the form

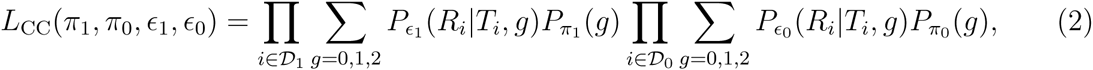

where *D*_1_ and *D*_0_ denote the sets of cases and controls, respectively, π*_d_* denotes the allele frequency for *D* = *d*, and (π_1_,*ϵ*_1_) and (π_0_,*ϵ*_0_) are separate parameters for cases and controls. Note that in writing (2) we assume that the depth *T* is independent of the genotype *G*. Also note that this formulation obviates the need to model other covariates (e.g., age and environmental exposures) as long as they are not confounders. The null hypothesis of the association test is *H*_0_ : *π*_1_ = *π*_0_. We re-parameterize (π_1_,π_0_) in terms of (*α*, *β*,) such that 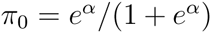 and 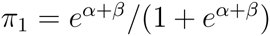; then the null hypothesis is *H*_0_ : *β=0*. The score statistic for *β* obtained from (2) is

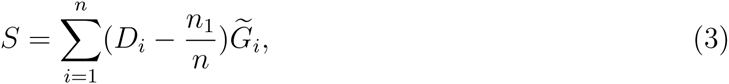

where

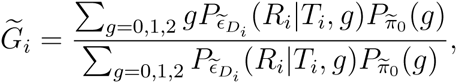

*n*_1_ is the number of cases, and (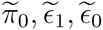) are restricted maximum likelihood estimates (MLEs) under the null; these restricted MLEs can be obtained via the expectation-maximization (EM) algorithm described in Supplemental Methods. 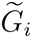 can be interpreted as the posterior dosage of the minor allele; as the read depth increases, 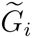 converges to the underlying genotype *G*_*i*_and *S* reduces to the standard score statistic 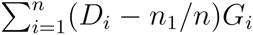. Finally, we construct the burden statistic *W* as a (weighted) sum of the score statistics at a set of SNVs in the gene of interest. The variance estimator *V* for *W* is calculated as the empirical variance of the efficient score functions (Lin 2006). When true SNVs are used, the test statistic 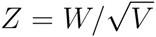 is asymptotically normal with mean 0 and variance 1.

The score statistic of the Derkach test (Derkach et al. 2014) has the same form as (3), as it also uses the posterior dosage 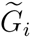. The only difference is that the Derkach test substitutes the genotype likelihood 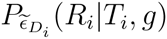 that is provided in the output of standard genotype calling packages (Li et al. 2009; DePristo et al. 2011), which calculate error rates based on *phred* scores.

### Screening out uninformative loci

In reality, the locations of rare SNVs are not available without calling genotypes. In order to include the maximum set of variants in the burden test without calling genotypes, we develop a screening algorithm to screen every locus (i.e., base pair) in the genome and filter out only loci that are “uninformative” in the sense that they yield *S* = 0 and thus do not contribute to the test statistic. Specifically, we consider the likelihood 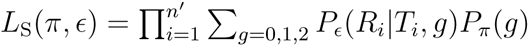 which is based on a homogenous group (i.e., cases or controls only) of *n′* subjects. Let 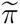 be the MLE based on *L_S_*(*π*,*ϵ*) under the constraint that *π* ϵ [0,1] and note that 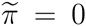 indicates no mutation in this group at this locus. Fortunately, we can easily determine whether 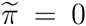 without iteratively solving for 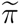. By definition, 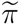 also maximizes the profile likelihood 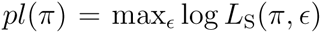. Because we have shown in Supplemental Methods that *pl*(π) is a concave function of π, a negative derivative of *pl*(π) at π = 0 leads to 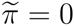. At π = 0, the *ϵ* maximizing log *L*_S_(π, *ϵ*) can be easily determined because, in the absence of any minor alleles, all reads carrying the minor allele must be errors. Therefore, we check the sign of *pl*(0) for cases and controls separately and screen out the loci at which both signs are negative. If 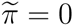 in both cases and controls, then 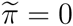 in the combined sample, where 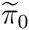 was defined in the text following expression (3). From 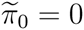, we have *G_i_* = 0 for all individuals and thus *S* = 0. This screening algorithm only involves evaluating simple (derivative) functions twice at each locus without any iteration, and is thus computationally extremely efficient.

### Bootstrap

Although most monomorphic loci are “uninformative” and will be screened out, there are exceptions. It is possible that a truly monomorphic locus has 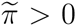 in one disease group or both, if by chance some individuals have more errors than expected. If a truly monomorphic locus has 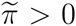 in the control group but 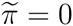 in the case group, the score statistic *S* of this locus will have a negative mean. Such loci will accumulate over the gene when controls have systematically lower depth (or higher error rate) than cases, and then the expected value of the burden statistic *W* will be substantially biased below zero, even when allele frequencies are identical among cases and controls at true SNVs. Consequently, screening for SNVs in the presence of differential sequencing qualities between cases and controls will invalidate the asymptotic version of our test.

We thus propose a bootstrap procedure for assessing the significance of the observed test statistic *Z*. The idea is to generate bootstrap datasets that mimic the original data in terms of read depth and error rate, have the same number of truly monomorphic loci and true SNVs, but have no difference in allele frequencies among cases and controls. To this end, we condition on the observed depth *T* and generate the minor-allele read count *R* using the estimated error rates 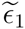 and 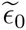 once the underlying genotype *G* is simulated. However, it is nontrivial to simulate *G*, because we do not know how many loci in the gene are true SNVs and what are allele frequencies at these SNVs. To obtain this information, we first form a “monomorphic” dataset by generating *R* at every locus in the gene assuming that all *G*s are zero; thus, each read for the minor allele is an error that occurs with rate 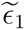 or 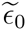, depending on the disease status. This dataset should provide a good approximation to the truly monomorphic loci in the original data, as the proportion of true SNVs in the original data should be small. Let *M_s_* be the number of loci that are screened in from the original data and let *F_s_*(*π*) be the cumulative distribution function (CDF) of estimated MAFs at the *M_s_* loci. Let M*_m_* and *F_m_*(π) be their counterparts in the monomorphic dataset. The CDF of allele frequencies at true SNVs, denoted by *F_p_*(*π*), is related to *F_s_*(*π*) and *F_m_*(*π*) through the equation

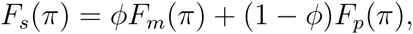

where *φ* is the proportion of monomorphic loci among loci that are screened in. This equation expresses the fact that the distribution of observed (non-zero) allele frequencies *F_s_*(*π*) in the original data is a mixture of the distributions for allele frequencies of true SNVs *F_p_*(*π*) and artifactual SNVs *F_m_*(*π*) that actually correspond to monomorphic loci. We estimate *ø* by 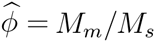 and *F_p_* by 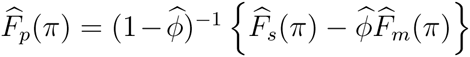, where 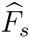 and 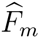 are empirical CDF estimators of *F_s_*(*π*) and *F_m_*(*π*) respectively. To ensure that 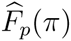 is monotonically increasing, we refine 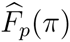 by fitting an isotonic regression to data points of 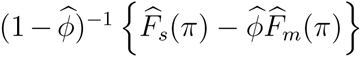 evaluated at the pooled (*M_s_* + *M_m_*) MAFs by the pooled-adjacent-violator algorithm (PAVA) (Robertson et al. 1988). After the largest value of MAF, we set 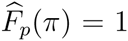. Finally, starting from the monomorphic dataset, we select 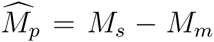 loci to be SNVs, sample *π* from 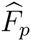, and re-generate *G* and *R* at these SNVs to form a final bootstrap dataset. Note that, for a small *π*, we may need to resample *G* repeatedly until each truly polymorphic locus screens in. The bootstrap statistic is then calculated based on all the loci that were screened in from the final bootstrap dataset. The entire procedure is repeated to generate multiple bootstrap replicates.

Although bootstrap tests are computationally intensive in general, we can save considerable time by adopting a sequential stopping rule (Besag and Clifford 1991). We stop after generating *L*_min_ bootstrap replicates, if these early replicates suggest a large *p*-value. When *L*_min_ = 5, the number of replicates at termination has a median of only 10 for a gene having no SNVs that affect the trait. We also use a closed sampling scheme, in which we restrict the total number of bootstrap replicates to be at most *K*_max_. If we stop when *L*_min_ bootstrap statistics exceed the observed *Z* and *K*_obs_ (≤ *K*_max_) replicates have been collected, we set the *p*-value to *L*_min_/*K*_obs_. If we stop when *K*_max_ replicates are reached and only *L*_obs_ (< *L*_min_) values exceed *Z*, we set the *p*-value to (*L*_obs_ + 1)/(*K*_max_ + 1).

### Adjusted empirical Bayes estimator for error rate

The MLEs of error rates may not recover the true distribution of error rates, which is essential for generating valid bootstrap replicates. In particular, when the true error rates are very small (e.g., ~0.02%), the MLEs tend to be over-dispersed. Therefore, we propose to use the empirical Bayes (EB) estimator in bootstrap instead of the MLE. We assume a prior beta distribution for error rates, i.e., *ϵ_j_* ~ *Beta*(*a*, *b*), where *j* = 1, …, *M*, *M* is the total number of loci in the gene, and *a* and *b* are hyperparameters that can be estimated by the method of moments (see Supplemental Methods). We show in Supplemental Methods that the EB estimator for *ϵ_j_* is 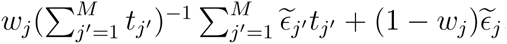, where 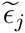 is the MLE, ω*_j_* = (*a* + *b*)/(*a* + *b* + *t_j_*), and *t_j_* is the total number of reads at locus *j* across all individuals. Thus the EB estimator imposes a shrinkage effect over the individual MLEs towards their (weighted) mean and alters the ranks of the MLEs according to *t_j_*. Additionally, because the EB estimator is known to over-shrink the distribution, we further adjust the EB estimates to be the quantiles of the prior beta distribution (Louis and Shen 1999) with estimates *a* and *b*. As long as *a* and *b* are consistently estimated, the true distribution can be accurately recovered.

### Read-based QC procedure

We have observed that a small proportion of read data (*R*, *T*) do not fit the binomial model (1). This may be due to genotype mosaicism (i.e., the presence of two or more populations of cells with different genotypes in one individual), experimental artifacts, sample contamination, or copy number variants. To detect data that do not fit the binomial model, for each individual at each locus that screens in, we calculated a likelihood-ratio-type statistic for the goodness of fit to the binomial model

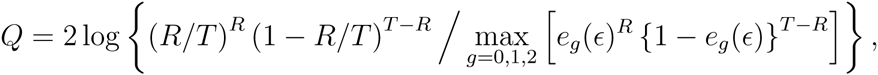

where *e_g_*(*ϵ*) = *ϵ*, 0.5, and 1 – *ϵ* for *g* = 0, 1, and 2, respectively. Then, we mask an individual at a variant (by setting *T* and *R* to zero) if *Q* is greater than 10 and remove a variant altogether if more than 5 individuals are masked at that locus. We can also identify individuals with problematic data by checking for the presence of an excessive number of *Q*s greater than 10.

## Results

### Simulation studies

We carried out extensive simulation studies to evaluate the performance of our proposed methods in realistic settings. We used the coalescent simulator cosi (Schaffner et al. 2005) to generate a base population of 100,000 European haplotypes with length 10 kb. We assumed that the 10 kb region corresponds to a gene with 3 exons that are separated by 2 introns, with introns being 3 times the length of exons. This setup gave us a total of 2,730 loci in exons, among which there are 44 SNVs with MAFs ≤ 0.05 in the base population. To generate individual genotypes, we sampled from the 100,000 haplotypes allowing recombination in introns (but not in exons). To generate disease outcomes, we considered a risk model that assumed equal attributable risk (AR) for each SNV: 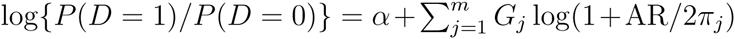, where *m* is the total number of SNVs, *G_j_* and *π_j_* are the genotype and MAF of the *j*th SNV, and *α* was set to –3 to achieve a disease rate of ~ 5%. This risk model implies that a more rare SNV has a stronger effect than a less rare SNV. The process was repeated until 500 cases and 500 controls were collected.

The sequencing reads *T* and *R* were generated to mimic real NGS data. We considered average read depths of 6×, 10×, and 30×, and average error rates of 0.02% and 0.016% (as observed in the UK10K cases and controls, respectively). While these very low error rates are characteristic of the newest Illumina platforms, we also considered average error rates of 1% and 0.5% that exist in historical NGS data (Nielsen et al. 2011). We sampled the locus-specific error rate *ϵ* from a beta distribution that yields the pre-specified average rate. We sampled the individual depth *T* by a two-step strategy which first simulates the locus-specific mean depth *c* from a beta distribution (re-scaled to achieve the pre-specified average depth) and then simulates individual *T*’s from a negative-binomial distribution with mean *c*. The first step permits the accessibility of sequencing to depend on local nucleotides, and the second step allows for dispersion in the individual count data. For specific parameter values in these distributions, refer to Supplemental Methods. Note that at each locus we sampled *ϵ* and *c* independently for cases and controls, even when the average values are the same between the two groups. Finally, we sampled *R* given (*T*, *G*, *ϵ*) according to (1).

We considered eight methods. First, we assumed that the 44 SNV locations were known and applied the asymptotic version of our method, the method using called genotypes that extends the multi-sample, single-locus genotyper seqEM (Martin et al. 2010) to allow for different error rates in cases and controls, the Derkach method using genotype dosages, and the method using true genotypes as a gold standard; we refer to them as New, CG, Dose, and True. Note that, to ensure fair comparisons, we used the error rates from our method in the implementation of the Derkach test, whose score statistic is then the same as our *S* in (3). Thus, although Derkach *et al*. used a slightly different variance estimator for the score statistic, New and Dose are asymptotically equivalent. Next, we considered the more realistic case that the SNV locations are unknown. We applied our method including the screening and bootstrap procedures and refer to it as New-SB. While this method aims to maximize the set of true SNVs, it may also include a sizable number of monomorphic loci that can adversely affect the power of association testing. We thus explored a modification of New-SB, which adds a thresholding step that excludes loci with estimated MAFs < (2*n*)^-1^ and is referred to as New-STB. The threshold of (2*n*)^-1^ corresponds to the MAF of a singleton variant and can effectively remove the majority of monomorphic loci that accidentally pass the screening algorithm, although at a cost of potentially losing some true singletons. In addition, we applied the method of called genotypes and the Derkach method based on loci that were screened in and refer to them as CG-S and Dose-S.

We focused on the weighted burden test of SNVs with MAFs ≤ 5%, in which each SNV is inversely weighted by 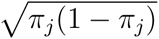 (Madsen and Browning 2009; Lin and Tang 2011); results of the unweighted test are provided in Supplemental Materials. We first evaluated type I error of the burden test using the aforementioned methods and summarized the results in Table 1. All of the new methods (New, New-SB, New-STB) have correct type I error, regardless of how different the sequencing depths and error rates are between cases and controls. The genotype calling methods (CG, CG-S) generally have inflated type I error when the average depths are different between cases and controls. Their type I error tends to be inflated even when the average depths are the same but there are random differences in individual regions between cases and controls; the inflation can be seen more clearly in Supplemental Table S1 which pertains to the unweighted test. Only when cases and controls have exactly the same sequencing feature at every locus, which can be achieved by sequencing cases and controls together, should the genotype calling methods have correct type I error (results not shown). The Derkach approach worked well when the SNV locations are known, but its type I error rate can be as much as 88 times the nominal level when the locations are unknown. In Table 2, we give additional results on the behavior of our test statistics under the null hypothesis. We see that the test statistic in the presence of screening is negatively biased from zero when controls have lower average depth than cases, which confirms the need for our bootstrap test. We also see in Table 2 that, when the average error rate is high, the screening procedure screened in a large number of monomorphic loci, and that the thresholding procedure effectively removed many such loci. Finally, we see that the bootstrap procedure accurately estimated the number of truly polymorphic loci. Supplemental Figure S1 shows that the MLEs of error rates are more dispersed than the true error rates (especially contain too many zeros when the average is 0.02%), the EB estimator imposed a strong shrinkage effect, and that our adjusted EB estimator accurately recovered the true distribution. Supplemental Figure S2 shows that, when the average error rate is 1%, the monomorphic loci that were screened in are typically associated with small 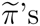, the majority of which are smaller than the threshold of (2*n*)^−1^.

**Table 1.** Type I error of the weighted burden test at the nominal significance level of 0.01.

| $c_1$ | $c_0$ | $\epsilon_1$ | $\epsilon_0$ | Known SNVs | | | | Unknown SNVs | | | |
| --- | --- | --- | --- | --- | --- | --- | --- | --- | --- | --- | --- |
|  |  |  |  | New | CG | Dose | True | New-SB | New-STB | CG-S | Dose-S |
| 6× | 6× | 0.02% | 0.02% | 0.010 | 0.011 | 0.009 | 0.009 | 0.011 | 0.011 | 0.011 | 0.009 |
| 30× | 6× | 0.02% | 0.02% | 0.010 | 0.055 | 0.009 | 0.009 | 0.010 | 0.010 | 0.033 | 0.161 |
| 30× | 30× | 0.02% | 0.02% | 0.009 | 0.010 | 0.009 | 0.010 | 0.010 | 0.010 | 0.010 | 0.010 |
| 30× | 6× | 0.02% | 0.016% | 0.011 | 0.061 | 0.010 | 0.011 | 0.009 | 0.009 | 0.029 | 0.143 |
| 10× | 10× | 1% | 1% | 0.008 | 0.010 | 0.008 | 0.009 | 0.011 | 0.008 | 0.012 | 0.011 |
| 30× | 10× | 1% | 1% | 0.008 | 0.037 | 0.008 | 0.010 | 0.011 | 0.008 | 0.358 | 0.878 |
| 30× | 30× | 1% | 1% | 0.010 | 0.011 | 0.011 | 0.010 | 0.011 | 0.009 | 0.012 | 0.012 |
| 30× | 10× | 1% | 0.5% | 0.011 | 0.024 | 0.011 | 0.010 | 0.011 | 0.008 | 0.379 | 0.702 |

$c_1$ and $c_0$ are average depths in cases and controls, respectively. $\epsilon_1$ and $\epsilon_0$ are average error rates in cases and controls, respectively. New: asymptotic version of our method. CG: method using called genotypes. Dose: the Derkach method using genotype dosage. True: method using true genotypes. New-SB: our method including the screening and bootstrap procedures. New-STB: our method including the screening, thresholding, and bootstrap procedures. CG-S: method using called genotypes at loci that were screened in by our screening algorithm. Does-S: the Derkach method using genotype dosage at loci that were screened in. Each entry is based on 10,000 replicates.

**Table 2.**
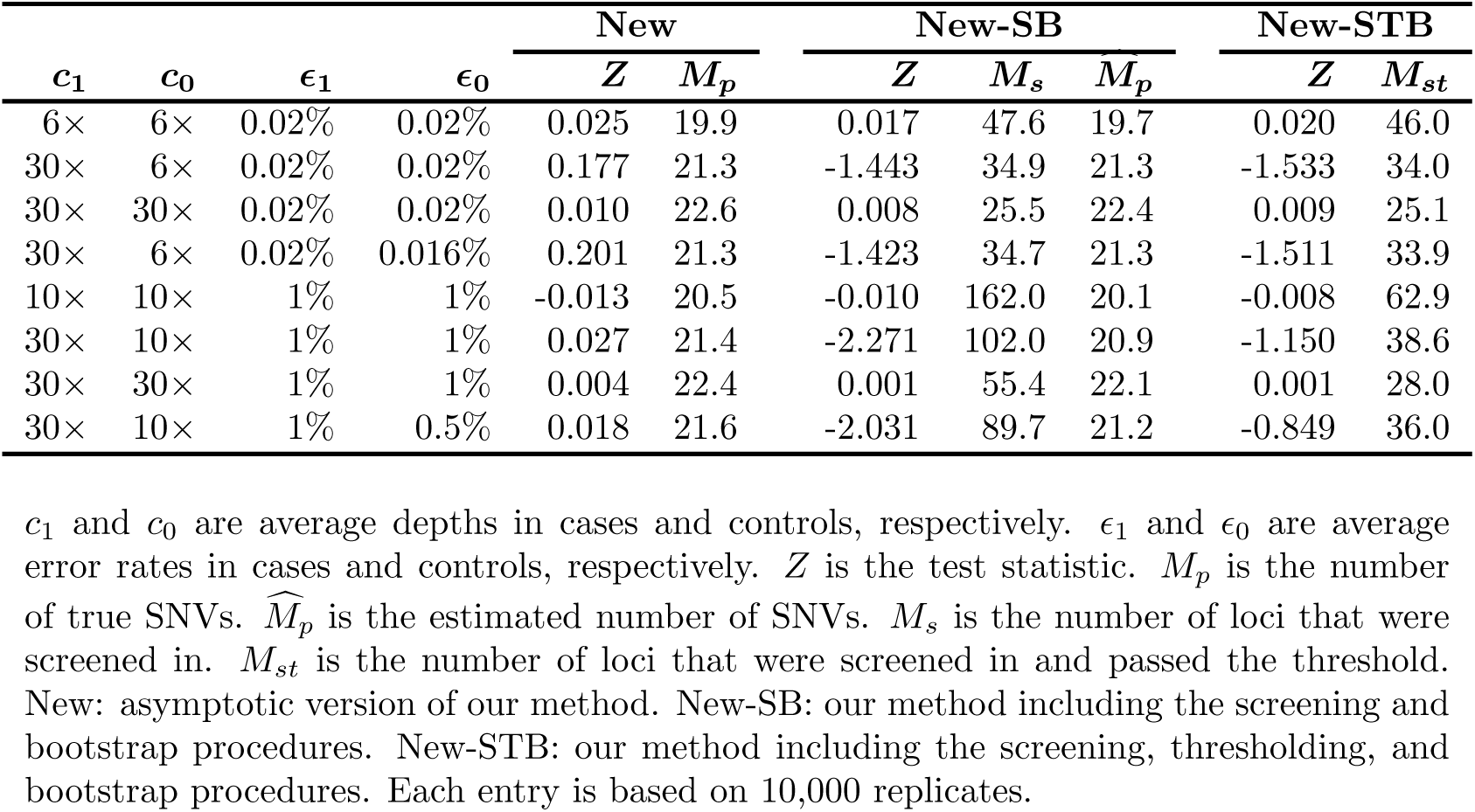
Other simulation results for the weighted burden test under the null hypothesis.

Figure 2 contrasts the power of different methods. The thresholding strategy implemented in New-STB significantly improved the power of New-SB at error rate of ~1% and performed as well as New-SB at ~0.02%. In the presence of differential depths between cases and controls, the power of CG-S and Dose-S can even decrease as the effect size starts to increase from zero and both are substantially lower than the power of New-SB and New-STB at median and high effect sizes. In the presence of equal average depths, the power of CG-S and Dose-S are comparable to that of New-SB and New-STB at error rate of ~0.02% and noticeably lower at ~1% (even at high depth of ~30×). Power curves pertaining to unweighted burden tests are displayed in Supplemental Figure S3, which shows similar patterns to Figure 2 but generally lower power due to the weighted nature of our simulation setup.

**Figure 2.**
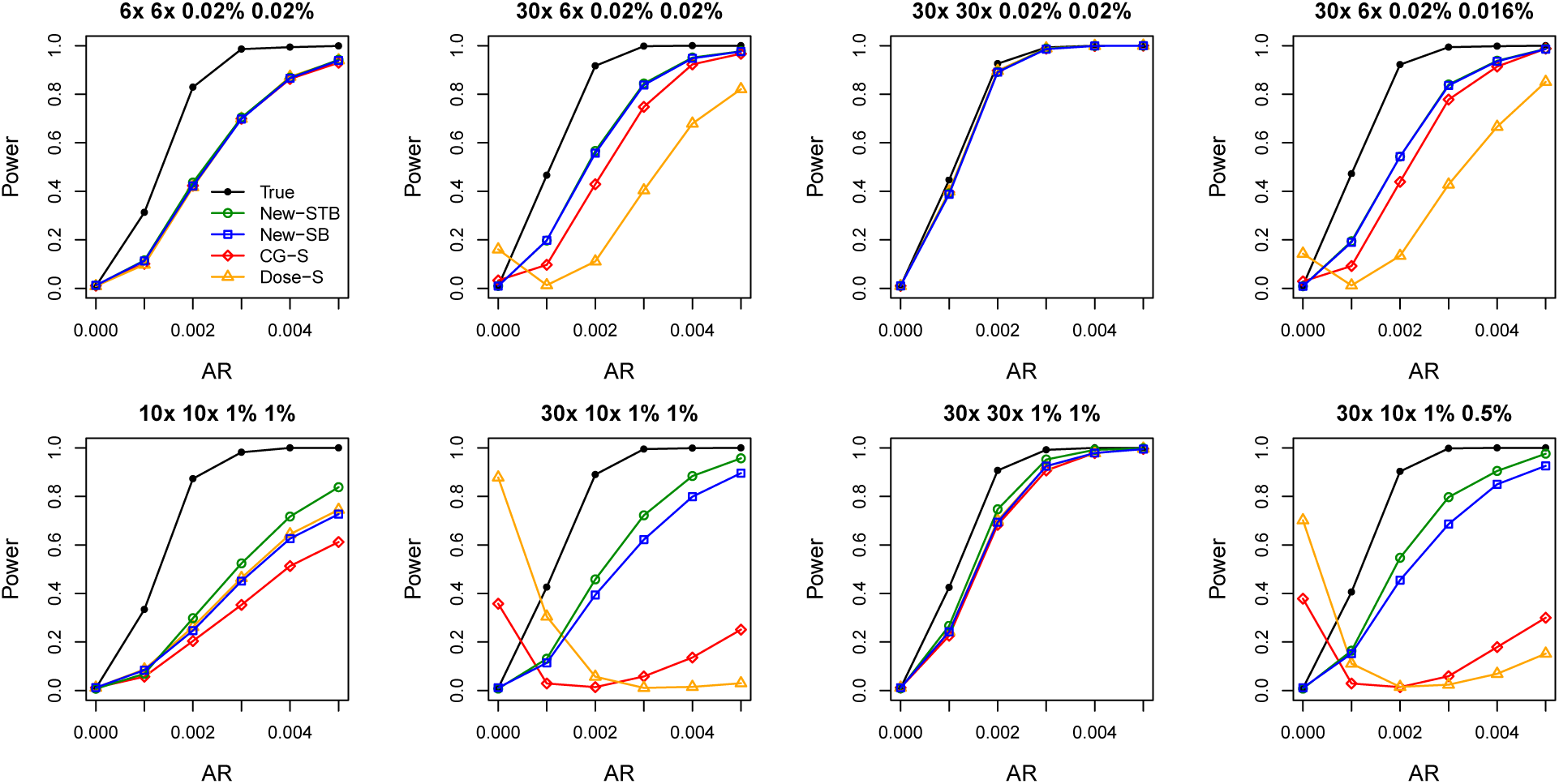
Power of the weighted burden test at the nominal significance level of 0.01. AR is the attributable risk per SNV. True: method using true genotypes. New-SB: our method including the screening and bootstrap procedures. New-STB: our method including the screening, thresholding, and bootstrap procedures. CG-S: method using called genotypes at loci that were screened in by our screening algorithm. Does-S: method using genotype dosage at loci that were screened in. Each power estimate is based on 1,000 replicates.

### UK10K data

The UK10K project was funded by the Wellcome Trust Sanger Institute in 2010 to help investigators better understand the link between low-frequency and rare genetic changes and complex human diseases by applying NGS on 10,000 people in the United Kingdom (UK). We focused on the samples collected by the Severe Childhood Onset Obesity Project (SCOOP), all of whom have severe, early onset obesity (i.e., body mass index Standard Deviation Scores (Must and Anderson 2006) > 3 and obesity onset before the age of 10 years). For controls, we utilized the population-based cohort collected in the TwinsUK study (randomly excluding one twin from each twinship) from the Department of Twin Research and Genetic Epidemiology at King’s College London. Both cases and controls are UK-based populations and part of the UK10K project. While the cases were whole-exome sequenced at average depth of 60×, the controls were whole-genome sequenced at average depth of 6×.

We used SAMtools to generate the pileup files from the BAM files and extracted read count data, filtering out reads that are PCR duplicates, that have mapping score < 30, that have improperly mapped mates, or that have *phred* base-quality scores < 30. We restricted our analysis to the consensus coding sequence gene sets (Pruitt et al. 2009) and further masked repeat regions, regions covered by monomorphic read alleles, and regions not covered by any reads, resulting in a total of ~14 million loci genome wide. We recorded read count data for these loci such that, for example, a locus covered by 10 reads of allele A and 1 read of C was coded as A10C1. Read count datasets in this format are much more manageable than the BAM files; our formatted, zipped files required only 126 GB of disk space, compared to ~14 TB for the BAM files. We obtained data in this format for 784 cases and 1,669 controls. We found that 87 cases had excessive read data that do not fit the binomial model (i.e., *Q* > 10) and we excluded these subjects (plus 1 additional case which is possibly in the same batch as the 87 cases) from further analysis; see Supplemental Methods for more details. Thus the analysis described here was based on 696 cases and 1,669 controls.

We considered two versions for the weighted burden test, one including all variants and one including only variants that are annotated as “probably damaging” or “possibly damaging” by PolyPhen (Adzhubei et al. 2013). We applied our methods, New-SB and New-STB, to scan all genes for association with severe childhood onset obesity. We set *K*_max_ = 10, 000, 000, which is sufficient for detecting *p*-values that pass the exome-wide threshold, which is on the order of 10^−6^. The analysis of damaging variants took a total of 1,713 hours on an IBM HS22 machine or equivalently 8.6 hours on 200 such machines in a computing cluster. We also applied the genotype calling method (CG-S) and the Derkach method (Dose-S) as described in Simulation Studies. Further, we analyzed the genotypes in the VCF files downloaded from the UK10K website. These genotypes were called by SAMtools, filtered by GATK VQSR, and imputed by Beagle (Browning and Browning 2009), by the UK10K investigators with cases and controls being processed separately. We refer to this approach as CG-VCF.

We screened in a total of 474,508 loci, among which 465,967 (98.2%) loci passed our read-based QC procedure. The 465,967 loci span over 16,318 genes; 431,311 passed the threshold of (2*n*)^−1^, and 288,535 were estimated to be polymorphic. Considering damaging variants only, 238,753 loci were screened in and passed QC; 219,540 passed the threshold, and 143,822 were estimated to be polymorphic. Note that the CG-VCF analysis was based on the same set of 465,967 loci, although some of them had been called monomorphic and were thus not included in the VCF files. As a result, the CG-VCF analysis included 167,980 loci, of which 79,271 were predicted as damaging.

The quantile-quantile plots are displayed in Figure 3. The observed *p*-values for New-STB and New-SB agree very well with the global null hypothesis of no association (genomic control λ =1), except at the extreme right tails. By contrast, the observed *p*-values for Dose-S, CG-S, and CG-VCF show very early departures from the global null distribution, reflecting severe inflation of type I error. Figure 4 shows that the test statistics are negatively biased from zero, which explained the poor performance of Dose-S.

**Figure 3.**
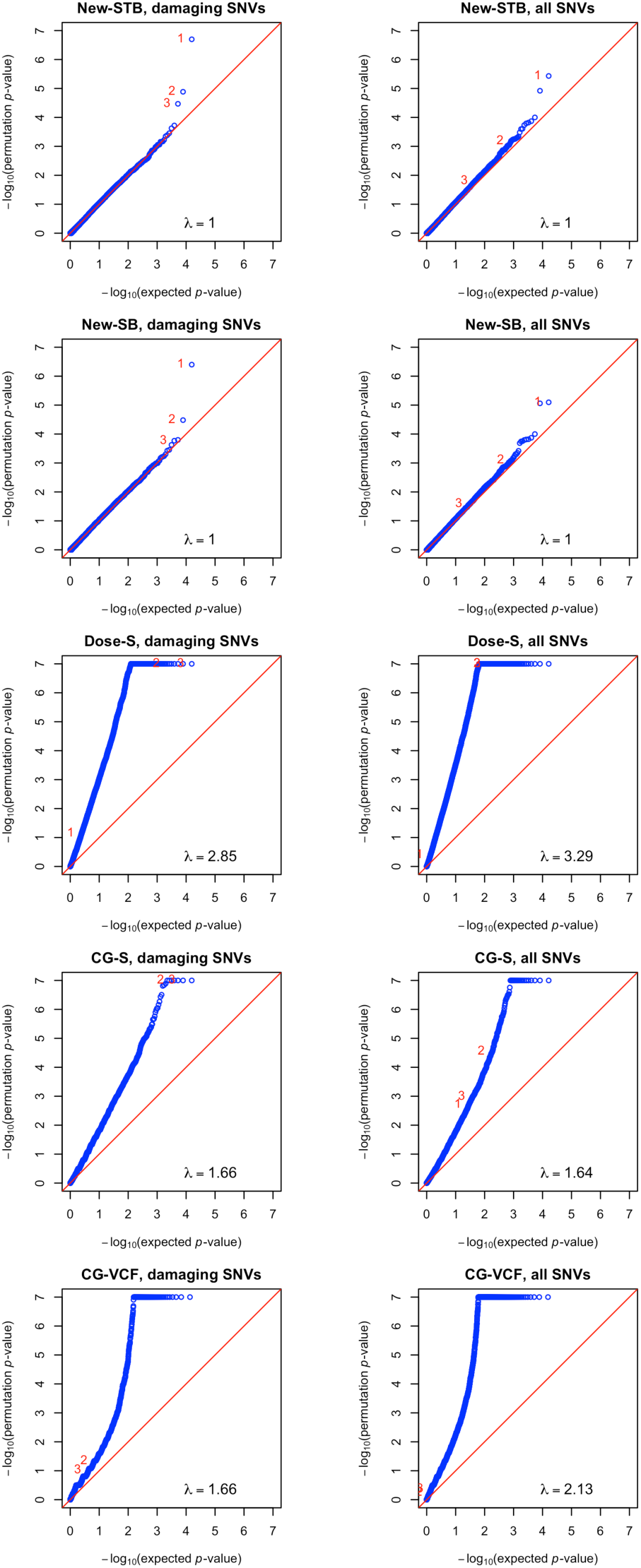
Quantile-quantile plots of —log_10_ (*p*-values) for the weighted burden test using damaging SNVs only (left side) and all SNVs (right side) in the analysis of the UK10K data. The top three genes identified by New-STB using damaging variants only are marked as 1–3. New-SB: our method including the screening and bootstrap procedures. New-STB: our method including the screening, thresholding, and bootstrap procedures. Does-S: the Derkach method using genotype dosage at loci that were screened in. CG-S: method using called genotypes at loci that were screened in. CG-VCF: method using genotypes in the downloaded VCF files

**Figure 4.**
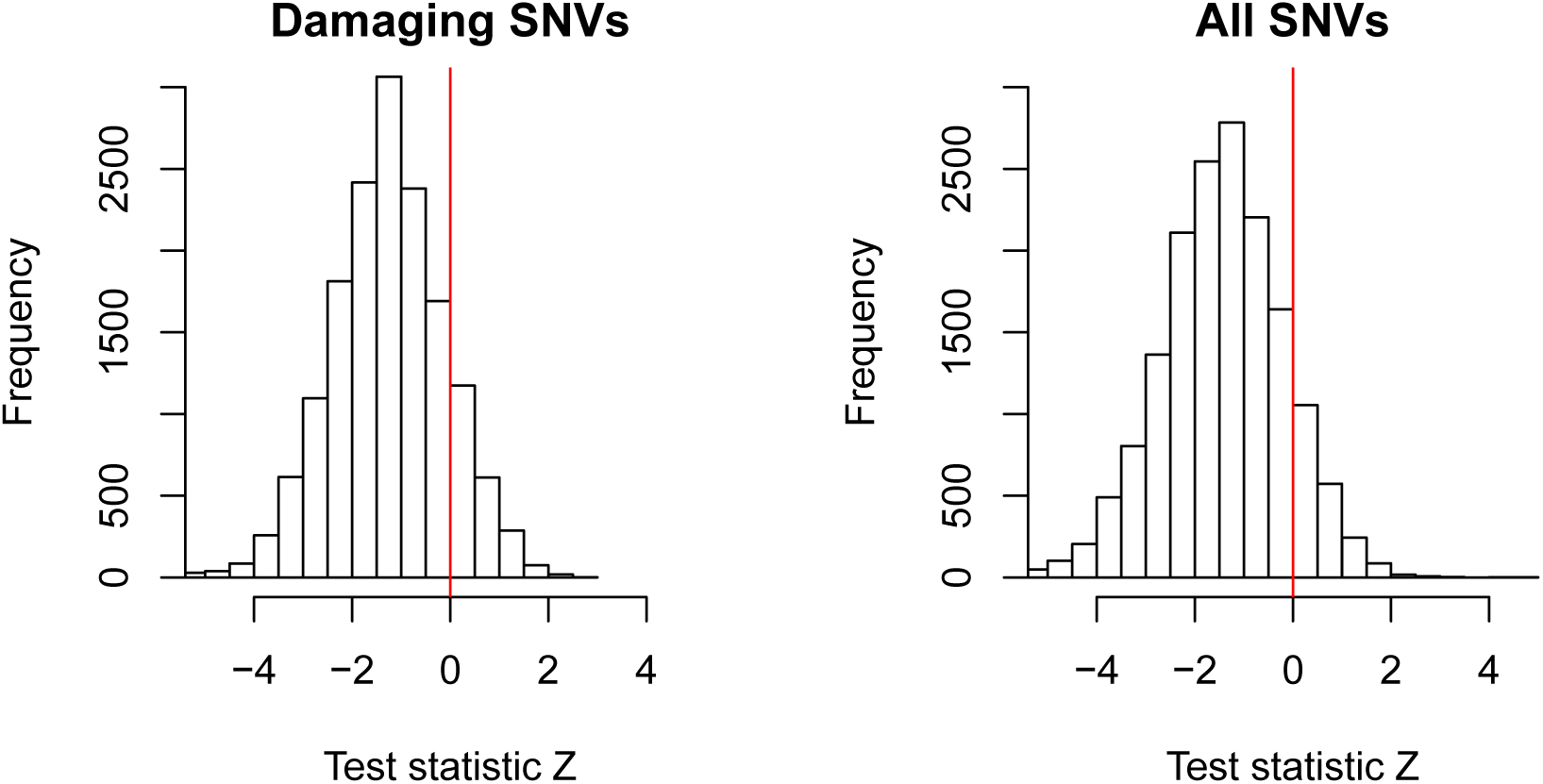
Distributions of the test statistic Z using damaging SNVs only (left side) and all SNVs (right side) in the analysis of the UK10K data. The left and right histograms are based on 15,659 and 16,318 genes, respectively.

Among all *p*-values generated by our methods, the smallest one, 2.0 × 10^−7^, was obtained for gene *BTBD18* by New-STB using damaging variants only, and this *p*-value passed the exome-wide significance threshold of 3.1 × 10^−6^ (0.05/16,318) after Bonferroni correction. Looking into the raw read data on this gene, we found that among cases the WES resulted in extremely low depth (~0.34×). (This kind of regions is not uncommon; indeed, 1.9% of all loci that were screened in have depth ≤1× in cases.) We found that at each of four loci (57512143, 57512745, 57513287, and 57513568 when mapped to the hg19 reference genome), there is a case individual covered by two reads and both are minor allele reads. These four suggestive minor allele homozygotes made large contributions to the score statistic and drove the gene-level association signal. As gene *BTBD18* has also been found to over-express in obese children elsewhere (NCBI GEO Profile ID: 64932244), it makes a plausible candidate for childhood onset obesity. Table 3 lists *BTBD18* and other top ten genes ranked by New-STB using damaging variants. Note that the standard genotype calling approach (CG-VCF) would have precluded *BTBD18* from association analysis due to the low depth data in cases. Using all SNVs, *BTBD18* was also ranked highest by New-STB (results not shown), with the same four loci driving the association signal, but the *p*-value did not pass the exome-wide significance threshold because of the inclusion of other neutral variants.

**Table 3.**
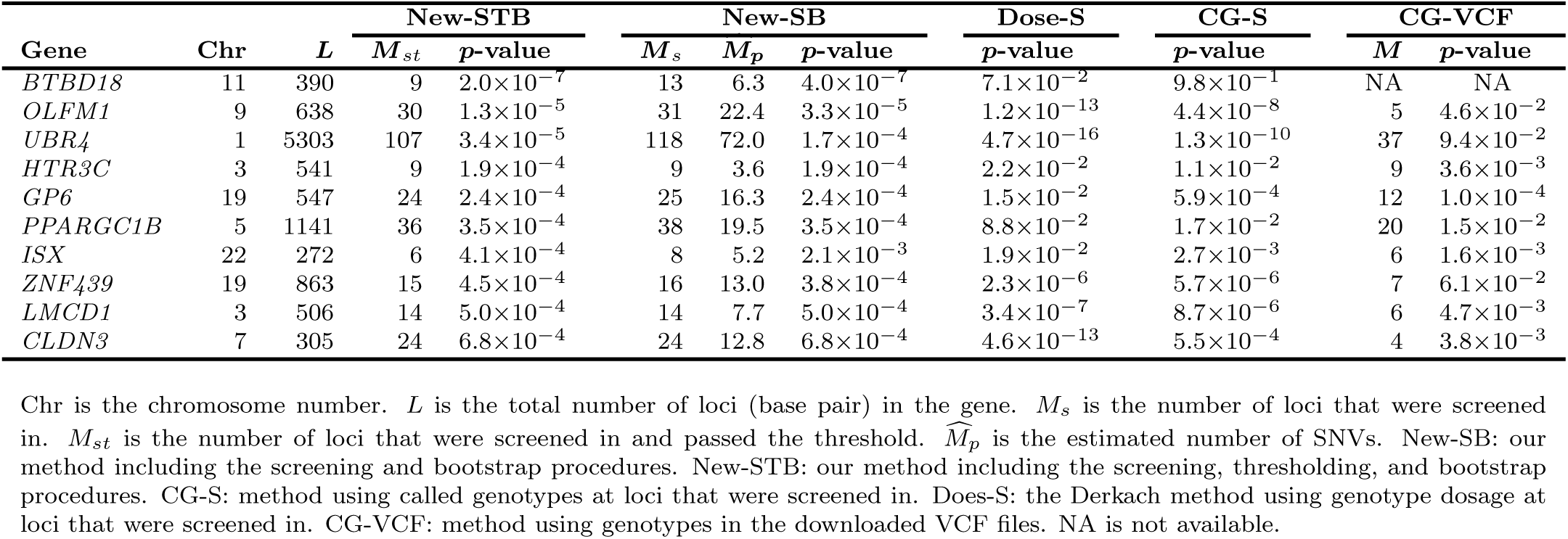
Top ten genes for childhood onset obesity identified by New-STB using damaging variants in the analysis of the UK10K data.

## Discussion

We have developed a robust and efficient approach to association testing of rare variants that is based on analyzing raw sequencing reads directly, without calling genotypes. It has been implemented in the publicly available software program TASER (Test of Association using SEquencing Reads). The robustness and efficiency of our methods make them ideal for the initial scan of association signals, where the precise genotypes are not of direct interest. Follow-up studies can be conducted to sequence the identified candidate genes at high depth, in which case the genotypes can be reliably identified and carefully examined for their functions.

Our read-based procedure allows use of far more loci than methods based on calling genotypes, because we do not filter out variants with low depth. For example, in analysis of the UK10K data, we only filtered out 1.8% of loci that were screened in; our final analysis included data from 465,967 loci. By contrast, the UK10K Statistics Group had to pare down to only 132,984 loci in order to achieve accurate type I error in the standard genotype calling approach (A Hendricks, pers. comm.), even though their analysis included almost 2,000 additional control participants from the Avon Longitudinal Study of Parents and Children (ALSPAC).

When developing our methods, we made some simplifying assumptions. First, we assumed independence (i.e., no LD) across rare variants when generating bootstrap replicates. This is reasonable because rare variants typically do not exhibit strong LD with each other (Pritchard 2001; Pritchard and Cox 2002). However, if strong LD occurs, it is possible to generate SNVs that have the same amount of LD as the original data by sampling haplotypes instead of single SNVs. The SNVs in the bootstrap sample can be placed in the same order (by allele frequency) as the original data.

Second, we assumed that base-calling errors are independent across loci. In reality, the base-calling errors might be correlated due to factors such as library preparation and sequence context. However, this assumption only affects the efficiency of our method, not its validity. We also assumed that the errors are symmetric, i.e., the probability of a read for the major allele being mis-called as the minor allele is the same as the probability of the minor allele being mis-called as the major allele. For analyzing rare variant data, this assumption has a negligible effect as rare variant homozygotes are extremely rare. Further, our methods estimate error rates directly from the read data, and thus ignored *phred* scores that characterize the base-calling quality and alignment scores that calibrate alignment quality. In our analysis of the UK10K data, we filtered out reads with alignment scores < 30 and *phred* scores < 30. We have shown in other work (unpublished) that *phred* scores and low-score reads can provide additional information. It would be possible to include a model of the variability in error rates that is explained by base-calling and alignment quality scores in our current approach.

We also assumed that all variants are biallelic. This assumption is reasonable because only a small fraction of SNVs have been verified to carry three or more alleles (Hodgkinson and Eyre-Walker 2010). In analyzing the UK10K data, we deleted in advance all calls for bases that differed from the two most frequent bases at every locus.

Finally, we do not account for confounders such as principal components for ancestry. In the UK10K data, all samples are UK-based Caucasians and are therefore not expected to have strong population stratification. It is also possible to extend our methods to allow confounders, by generating bootstrap replicates that have the same amount of confounding as the original data. We plan to describe such approaches in a subsequent report.

We have focused on the burden test in this article. Because our score statistic may not have mean zero after screening, it is nontrivial to construct the sequence kernel association test (SKAT) (Wu et al. 2011). A valid SKAT statistic requires the score statistic be properly centered; we are currently developing methods to center the score statistic within our bootstrap approach.

There is resurgent interest in generating NGS data for case-parent trio studies. It may occur that individuals within a family are sequenced with different depths and error rates. For example, the affected child may be sequenced with a technology that produces higher depth and fewer errors. The standard transmission disequilibrium test (TDT) (Spielman et al. 1993) and its extensions to rare variants (De et al. 2013) are very sensitive to genotyping errors and tend to yield inflated type I error in the presence of sequencing differences within trios. We are currently extending our read-based framework to trio studies.

In summary, we have developed a tool to perform association testing of rare variants that allows the case and control samples to be sequenced using different platforms. We showed that the proposed approach has correct type I error under various practical scenarios and is more powerful than the genotype calling approach when the latter is valid. Using the UK10K data, we demonstrated that the proposed approach has the potential to discover rare variants that are associated with complex human traits.

## Data access

The URL for the software TASER is **http://web1.sph.emory.edu/users/yhu30/software.html**

## Acknowledgments

This study makes use of data generated by the UK10K Consortium, derived from samples from the Severe Childhood Onset Obesity Project (SCOOP) and the TwinsUK study. A full list of the investigators who contributed to the generation of the data is available from **www.UK10K.org**. Funding for UK10K was provided by the Wellcome Trust under award WT091310. This research was supported in part by the University Research Committee of Emory and the National Institute of General Medical Sciences of the National Institutes of Health under Award Number R01GM116065–01A1.

## Disclaimer

The findings and conclusions in this report are those of the authors and do not necessarily represent the official position of the Centers for Disease Control and Prevention.

